# Wide-field calcium imaging of cortical activation and functional connectivity in externally- and internally-driven locomotion

**DOI:** 10.1101/2023.04.10.536261

**Authors:** Sarah L. West, Morgan L. Gerhart, Timothy J. Ebner

**Author notes:** Correspondence: Timothy J. Ebner, M.D., Ph.D. Department of Neuroscience, University of Minnesota, Lions Research Building, Room 421, 2001 Sixth Street S.E. Minneapolis, MN 55455. Competing Interests Statement: The authors declare no competing financial interests.

## Abstract

The neural dynamics underlying self-initiated versus sensory driven movements is central to understanding volitional action. Upstream motor cortices are associated with the generation of internally-driven movements over externally-driven. Here we directly compare cortical dynamics during internally- versus externally-driven locomotion using wide-field Ca^2+^ imaging. We find that secondary motor cortex (M2) plays a larger role in internally-driven spontaneous locomotion transitions, with increased M2 functional connectivity during starting and stopping than in the externally-driven, motorized treadmill locomotion. This is not the case in steady-state walk. In addition, motorized treadmill and spontaneous locomotion are characterized by markedly different patterns of cortical activation and functional connectivity at the different behavior periods. Furthermore, the patterns of fluorescence activation and connectivity are uncorrelated. These experiments reveal widespread and striking differences in the cortical control of internally- and externally-driven locomotion, with M2 playing a major role in the preparation and execution of the self-initiated state.

## Introduction

Externally-driven movements are movements directed by an outside stimulus, such as an external sensory cue. Internally-driven movements are, instead, self-directed or self-determined by the actor. Externally- and internally-driven movements are thought to utilize different cerebral cortical mechanisms. In both humans and non-human primates, the premotor and supplementary motor area (SMA) are associated with driving internally-driven movements ^1–3^. Conversely, the dorsal steam with its projections from sensory areas to parietal cortex and then to frontal motor areas is associated with externally-driven movements ^4, 5^. Therefore, internally- versus externally-driven actions are likely to generate different activation and functional connectivity (FC) patterns cortex-wide.

In previous experiments ^6^, we found that spontaneous locomotion exhibits a unique correlation pattern in the cortical functional network compared to rest. In this correlation pattern, most dorsal cortical areas decrease in correlation with other cortical areas, but the most anterior portion of M2 shows an increase in correlation with the rest of the cortex. Because of M2’s association with the control of internally-generated movement, we hypothesized that M2 FC is lower during externally-driven locomotion than internally-driven locomotion. We also hypothesized that patterns of cortical activation and FC would differ well beyond M2 and involve much of the cerebral cortex. To obtain this wider view of cerebral cortical engagement, we performed wide-field Ca^2+^ imaging on mice as they walked on a motorized treadmill and compared the activation and network FC of the dorsal cerebral cortex to that in spontaneous locomotion.

Using partial least squares regression (PLSR) and regressing out the contributions of potentially confounding motor and behavior parameters, such as arousal level and locomotion speed, we show marked differences in both the patterns of cortical activation and FC in motorized and spontaneous locomotion. During the starting and stopping periods, M2 exhibits much higher FC during spontaneous than during motorized locomotion. In contrast, motorized locomotion is characterized by much larger and more wide-spread increases in activation in the starting period compared to spontaneous locomotion. Overall FC is lower for motorized compared to spontaneous locomotion for all behavior periods and the decrease is particularly striking in M2, further highlighting M2’s differential roles in internally- and externally-driven locomotion. During the pre-starting period, there are limited FC changes for motorized locomotion, while spontaneous is characterized by developing global preparatory increases in FC in all but the anterior M2 nodes. Finally, fluorescence activation and FC are fundamentally different, providing two independent forms of information encoding. Therefore, not only does M2 have differential roles in internally- and externally-driven locomotion, but our results also emphasize that the spatial patterns of activation and FC across the dorsal cerebral cortex are fundamentally different.

## Results

### Wife-field Ca^2+^ fluorescence imaging in motorized and spontaneous locomotion

Seven Thy1-GCaMP6f transgenic mice were trained to walk head-fixed on a horizontal disk motorized treadmill through pseudo-randomly timed stages of rest, locomotion starting, steady-state walk, stopping, and changes in speed (see Methods; Figure 1A-C). These same mice were also allowed to walk or rest spontaneously immediately before or after motorized treadmill sessions. We collected a total of 62.5 hours of wide-field Ca^2+^ fluorescence imaging during the motorized task (mean 8.9 ± 2.0 hours per mouse), in addition to 30.8 hours during spontaneous locomotion (mean 4.4 ± 0.9 hours per mouse), which resulted in ∼1500 occurrences of each transition type. The total number of occurrences of each behavior period collected and analyzed after data cleaning steps (see Methods) is shown in Supplementary Table 1. Mice walked at an average speed of 7.0 ± 3.4 cm/s during spontaneous locomotion. We used spatial Independent Component Analysis (sICA) to find 32 functional regions consistent across mice, called “nodes,” falling within 6 approximate anatomical cortical regions, labeled here as secondary motor (M2), primary motor (M1), somatosensory (S1), lateral parietal (LP), mid-parietal and visual (MV), and retrosplenial (Rs) cortices (Figure 1D and E). Mean fluorescence time series (the percent change in fluorescence divided by baseline fluorescence, ΔF/F%, see Methods) were extracted from each node (Supplementary Figure 1), and the FC between nodes was calculated as the Pearson correlation of the fluorescence time series, resulting in a total of 496 unique node-node correlation pairs.

**Figure 1.**
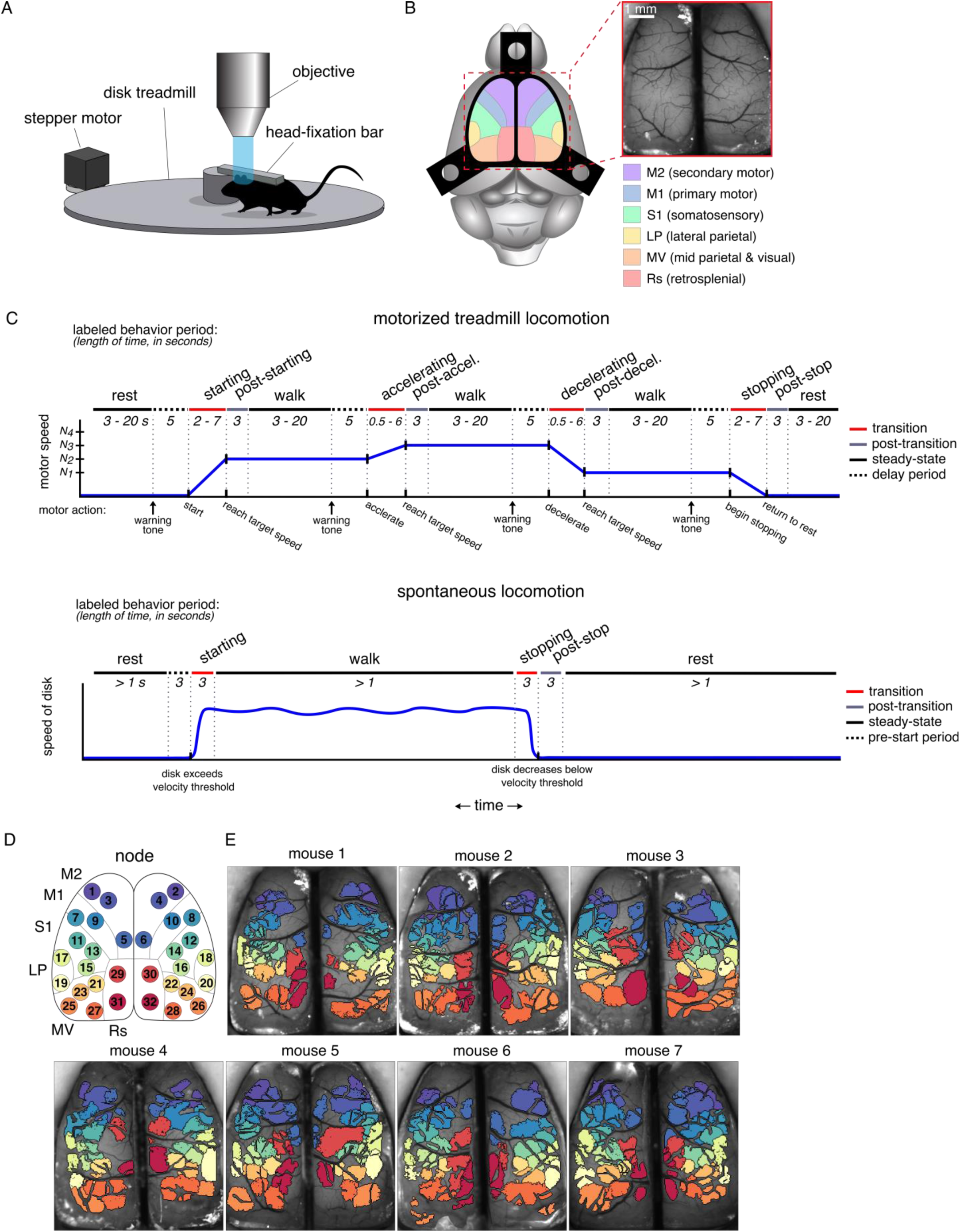
Motorized treadmill locomotion experimental setup. **A.** Experimental set-up for Ca^2+^ fluorescence imaging of head-fixed mice in the motorized treadmill task. Modified from West et al. (2022). **B.** Approximate cortical regions, as defined by the Allen Brain Atlas Common Coordinate Framework ^7^, observable through the polymer skull (inset) by epifluorescence microscopy. Scale bar = 1 mm. Common Framework colors correspond to cortical regions defined in key. Modified from West et al. (2022). **C.** Behavioral tasks are divided into behavioral periods for motorized locomotion (top) and spontaneous locomotion (bottom). **D.** Approximate cortical locations of the 32 nodes found common across all mice. **E.** The spatial independent component (IC) domains used for each mouse. The color of each IC corresponds to the assigned node ID, colors shown in D. Note that the blood vessels within an IC are removed, accounting for their fragmented appearance (see Methods).

### Cortical activation and functional connectivity in motorized and spontaneous locomotion

We first used PLSR to remove the effects of speed, acceleration rate, pupil diameter, and duration from the Ca^2+^ fluorescence activity and FC across the cortex in each behavior period (see Methods and Supplementary Figure 2). Then the differences in fluorescence activity and FC were determined across behavior periods and between internally- and externally- driven locomotion.

The neural activation patterns based on PLSR of the Ca^2+^ fluorescence reveal marked differences between motorized and spontaneous locomotion (Figure 2A and C) and agree with the time courses observed in the raw ΔF/F% (Supplementary Figure 1). Figure 2A and C display changes in activation of the different locomotion periods compared to rest, while Supplementary Figure 4A and C show changes across sequential behavior periods. Motorized locomotion is characterized by cortex-wide activation increases coincidental with locomotion onset (Figure 2Ai) that persists into the post-starting period (Figure 2Aii). Spontaneous starting also exhibits widespread increases, although they are approximately half the magnitude of those in motorized (Figure 2Ci). In both conditions, these activations diminish once steady-state walk is reached, during which nodes in S1 show the largest activations compared to rest (Figure 2Aiii and Cii). Interestingly, once stopping is initiated in the motorized condition, activations across the cortex drop to levels lower than those at rest, with the largest decreases occurring in peripheral nodes (Figure 2Aiv). Meanwhile, during spontaneous stopping the increases present at steady-state walk are maintained, with further increases in the most anterior nodes of M2 (nodes 1 and 2; Figure 2Ciii and Supplementary Figure 4Ciii). Therefore, both at beginning and end of locomotion, the changes in fluorescence across the dorsal cortex are markedly different for internally- versus externally- driven locomotion, consistent with distinct control processes and patterns of cortico-cortical communication.

**Figure 2.**
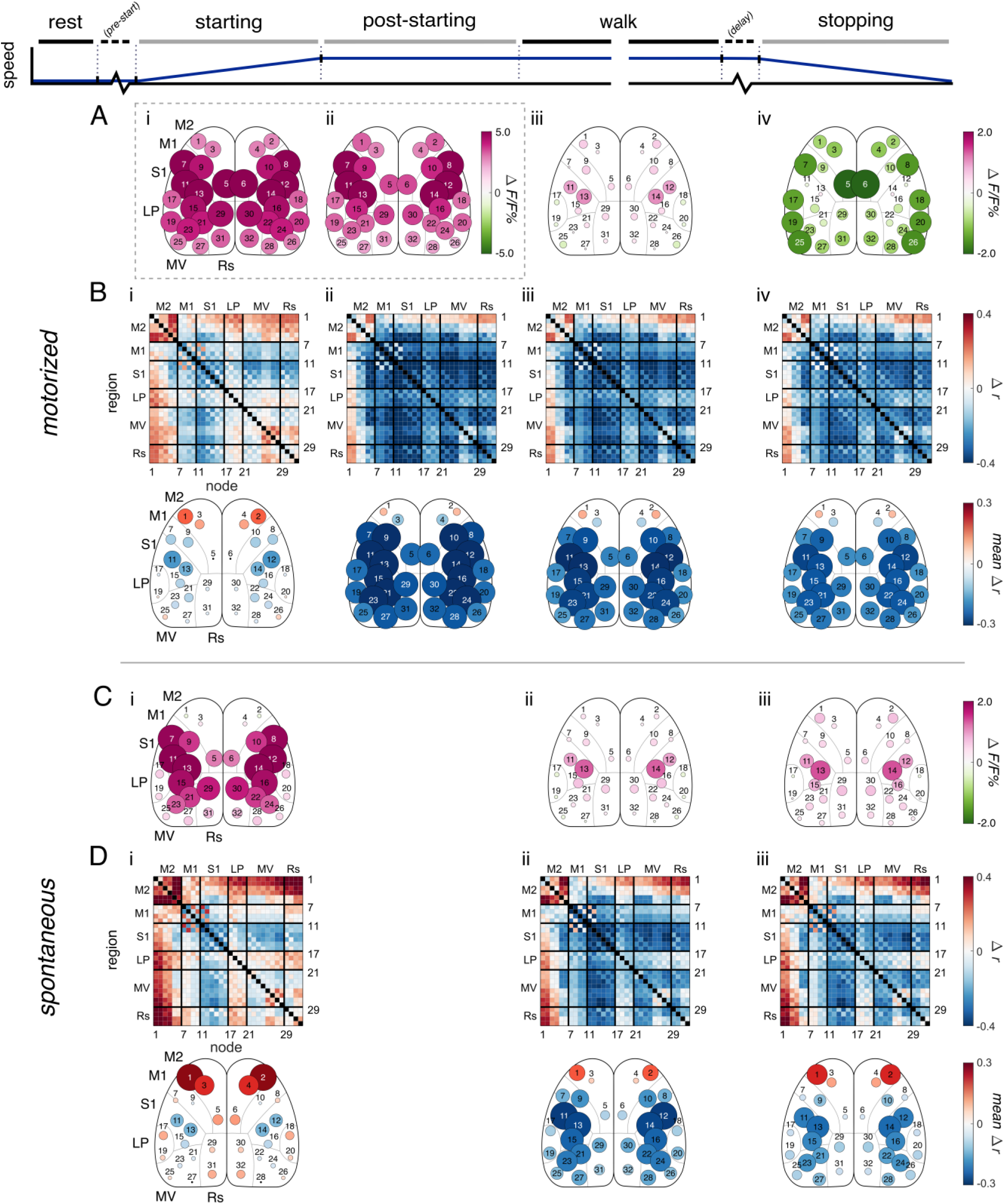
Change in correlation in motorized treadmill and spontaneous locomotion. **A.** Significant change in fluorescence activity of motorized treadmill behavior periods compared to the motorized treadmill rest period (α<0.05, permutation test with false discovery rate correction). Fluorescence is expressed as a % of the mean fluorescence for that node. **B.** (top) Significant change in correlation of motorized behavior periods compared to the motorized rest period (α<0.05, permutation test with false discovery rate correction). (bottom) Graph representation of the total mean changes in correlation with all other nodes for each node α<0.05, permutation test with false discovery rate correction). **C.** Similar to A, but for spontaneous locomotion during behavior periods. **D.** Similar to B, but for spontaneous locomotion behavior periods. Changes in fluorescence and correlations during pre-start, post-stop, and delay periods are not included (see Figure 4).

We next compared the FC of each behavior period in motorized and spontaneous locomotion to FC at rest. The baseline FC at rest for each condition are very similar (see Supplementary Figure 3), and each undergo a distinct change in the cortical functional network during locomotion (Figure 2B and D). Additional comparisons in FC between sequential behavior periods are provided in Supplementary Figure 4. FC decreases throughout much of the cortex for both spontaneous and motorized locomotion, except nodes in M2 (Figure 2B and D). These M2 nodes, particularly the most anterior (nodes 1 & 2), becomes more correlated with most of the cortex in locomotion starting (Figure 2Bi and Di). Notably, the spontaneous starting period shows the largest increase in M2 FC (Figure 2Di), while the motorized condition is characterized by larger FC decreases throughout locomotion (Figure 2B). Both conditions display a continuation of the steady-state walk FC pattern into the stopping period (Figure 2Biii and Diii). However, inspection of the specific changes from walk to stopping reveals small increases in FC throughout most of the cortex, with the notable exception of the decreases in anterior M2 FC in the motorized condition (Supplementary Figure 3Biv and Diii). In both conditions, the FC of LP nodes elevate modestly during starting (Figure 2Bi and Di). These LP nodes likely reflect the barrel cortex^7^. The changes in the FC of these parietal nodes may be due to the modulatory effects of locomotion on primary sensory cortices ^8, 9^, and the tendency we observed of the mice to begin whisking as they initiate walking.

Comparison of the modulation in Ca^2+^ fluorescence with the modulation of FC highlights the dissimilarity of these two metrics. The widespread and large changes in fluorescence during starting in the motorized condition are not reflected in the FC (Figure 2Ai versus 2Bi). Conversely, the widespread decreases in correlation throughout the cortex in both conditions during walk and stopping do not match the more muted and localized changes in fluorescence. To quantify the relationship between activation and FC, we calculated the correlation between the change in activation and the change in total connectivity across all analyses for each node (calculated for 16 pairs of contralateral nodes for the change across behavior periods, speed, acceleration rate, duration, and pupil diameter cases equaling a total of 80 relationships). Notably, changes in fluorescence and changes in FC are not significantly related in the vast majority of cases (74 of 80), with only a few exceptions (nodes 11 and 12 together show a negative correlation in changes occurring across behavior periods, nodes 19-24 and 29-32 show a positive correlation in changes as speed increases; t-test with False Discovery Rate correction; data not shown). Therefore, FC is not simply a reflection of neural activity. The results support that correlation and activation are distinct modes of information processing and representations of behavior.

### Differences in cortical activation and connectivity between motorized and spontaneous locomotion

We next directly compared fluorescence and FC between motorized and spontaneous locomotion in each behavior period (Figure 3). Major differences in Ca^2+^ fluorescence are apparent, with motorized locomotion characterized by large uniform increases in fluorescence for all nodes during starting (Figure 3Aii) and large uniform decreases during stopping compared to spontaneous (Figure 3Aiv). However, there is little difference between the two conditions during the steady-state rest and walk periods (Figure 3Ai & iii, respectively). As for FC, the motorized condition is generally lower than spontaneous in each period (Figure 3), but especially during the starting and stopping transition periods (Figure 3Bii & iv). Notably, the motorized condition also shows additional reduction in M2 FC in these transition periods (Figure 3Bii & iv). As described above, this occurs in the stopping period as M2 FC decreases from steady-state walk to stopping in the motorized condition but increases in spontaneous (Supplementary Figure 4Biv and Diii). Importantly, there is minimal difference in M2 FC between the two conditions during steady-state walk (Figure 3Biii). There is also increased FC in the motorized condition between contralateral M1 nodes, particularly in starting and walk (Figure 3Bii and iii), as well as between lateral M1 and the rest of the cortex in both steady-state rest and walk (Figure 3Bi and iii). Finally, FC increases between M2 and posterior MV nodes, as well as between M2 and Rs, in steady-state rest and walk (Figure 3Bi and iii). Overall, these results reveal that motorized and spontaneous locomotion, while similar during steady-state behavior, differ greatly in cortico-cortical engagement during transition periods, with spontaneous showing elevated anterior M2 nodes communication with the rest of the cerebral cortex.

**Figure 3.**
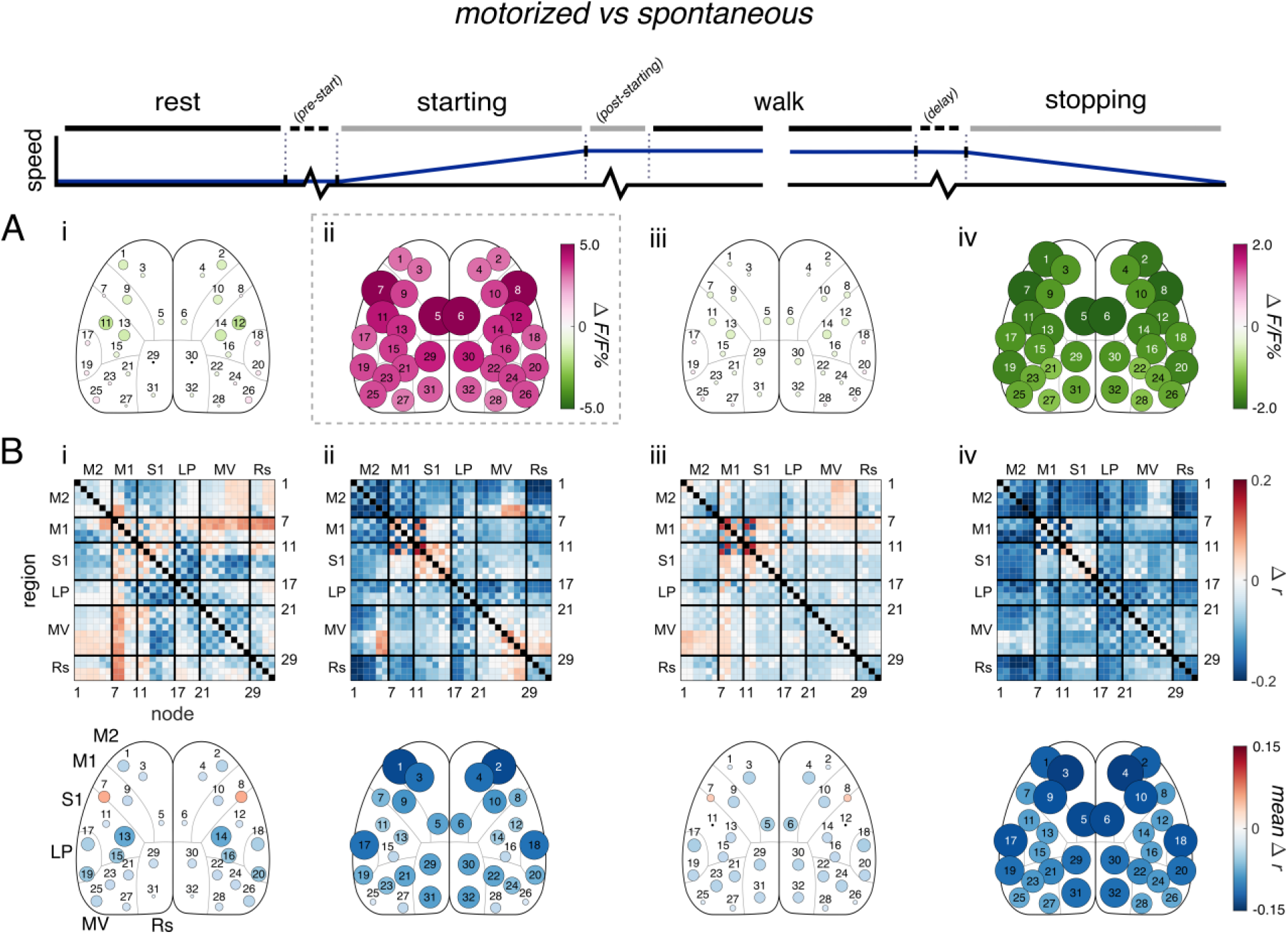
Motorized vs spontaneous locomotion. **A**. Significant changes in fluorescence across experimental conditions (from spontaneous to motorized treadmill locomotion, motorized minus spontaneous; α<0.05, permutation test with false discovery rate correction). Positive changes indicate higher fluorescence amplitude in the motorized treadmill condition at a given node. **B**. Similar to A, but for significant changes in FC. Positive changes indicate higher correlations in the motorized condition.

### Differences in pre-start and post-stop periods between motorized and spontaneous

To further investigate the differences between the motorized and spontaneous conditions in the locomotion onset and offset transition states, we examined cortical activation and FC during locomotion preparation and immediately after offset. We defined locomotion preparation as the 3 s immediately before locomotion onset (“pre-start” period; Figure 1C). In the motorized condition, auditory cues were given 5 s before the treadmill started, stopped, or changed speed, and the last 3 s of these windows were analyzed as preparatory periods (see Methods). As a control, specialized cues were given when the treadmill did not change speed (“control delay” periods, which preceded “maintaining” periods). The pre-start period in the motorized condition was compared to this control delay period to account for any effect of the warning tones on the cortical state, and the spontaneous pre-start period was compared to steady-state rest (Figure 4). The direct comparison between the pre-start periods is show in Supplementary Figure 5A.

**Figure 4.**
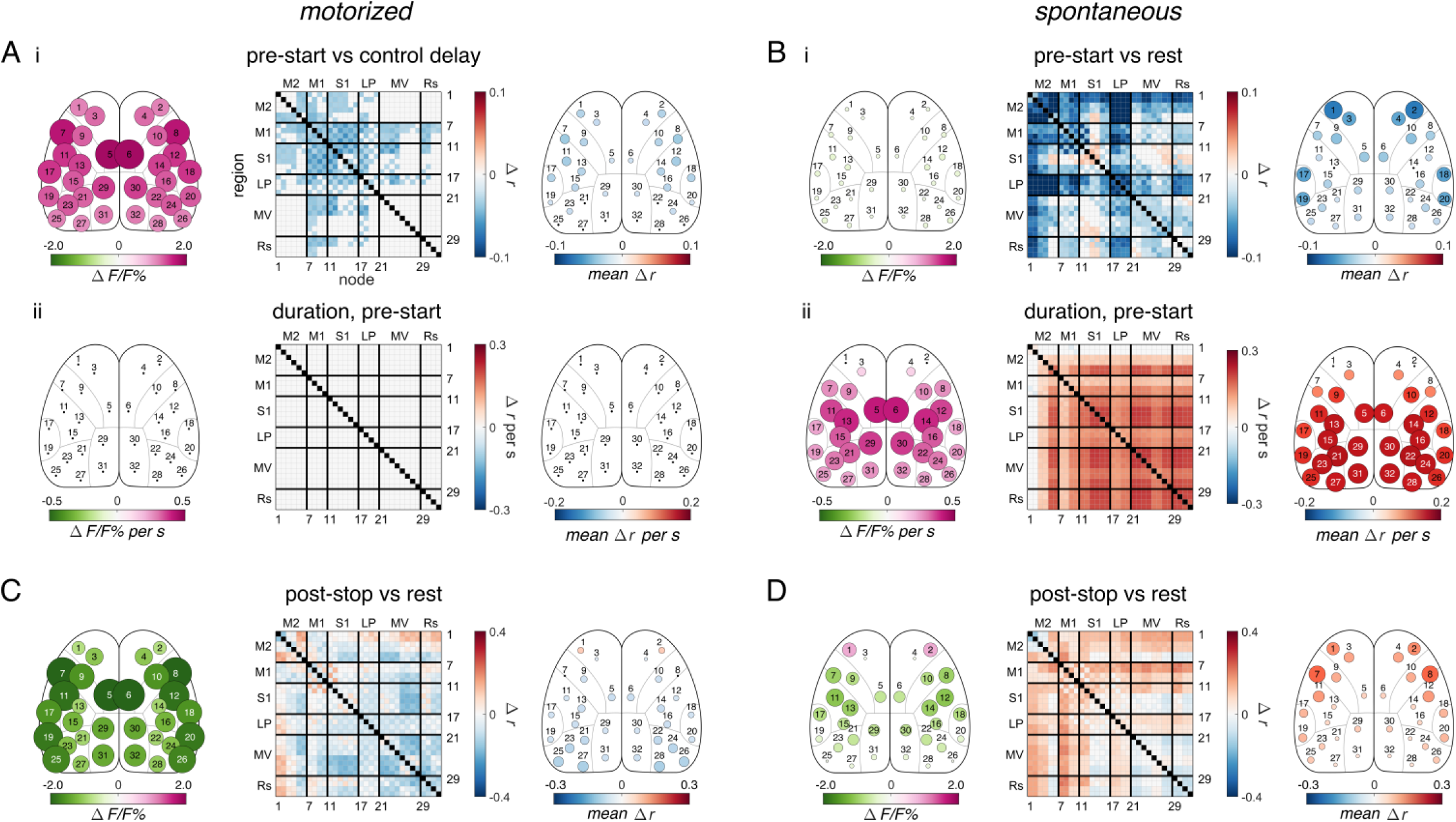
Differences in pre-start and post-stop between motorized treadmill and spontaneous locomotion. **A.** (i) Significant changes in fluorescence (*left*) and correlation (*middle and right*) in the motorized treadmill pre-start period compared to the control delay period as calculated and displayed in Figures 2 & 3. (ii) Significant changes in fluorescence (left) and correlations (right) with duration over the motorized pre-start period (note that no changes were significant). **B.** Similar to A, but for the spontaneous pre-start period, with (i) compared to spontaneous rest. **C.** (i) Significant changes in fluorescence (left) and correlation (middle and right) in the motorized post-stop period compared to rest. **D.** Similar to C, but for the spontaneous post-stop period.

Cortical activation and FC differ greatly between the motorized and spontaneous conditions during the pre-start period. Motorized locomotion displays large increases in activation in all nodes (Figure 4Ai), with only small decreases in FC, which are marginally strongest in M1 nodes (Figure 4Ai). Conversely, the pre-start period before spontaneous locomotion shows only marginal decreases in activation across the cortex, but the most anterior nodes of M2 exhibit marked decreases in FC (Figure 4Bi). The changes in activation and FC from the beginning of the pre-start period until the onset of locomotion were calculated as the results of the PLSR regression of duration against FC in the pre-start periods (see Methods). No changes in activation or FC occur over the duration of the pre-start period before motorized locomotion (Figure 4Aii), but large, global increases in FC occur in all but the anterior M2 nodes before spontaneous locomotion (Figure 4Bii). This agrees with previous findings that spontaneous locomotion is preceded by increases in activation and FC across the cortex^6^. There is also a drop in FC in LP nodes, which may correspond to the tendency of the mice to whisk before walking.

Similarly, we observe substantial differences between the motorized and spontaneous conditions after locomotion termination, defined as the 3 s immediately following offset (“post-stop” period; Figure 1C). In the motorized condition, there are large decreases in activation, particularly in lateral M1 and medial M2 nodes, and the locomotion FC pattern is still partially present (Figure 4C). The spontaneous condition shows a unique increase in anterior M2 activation as well as an increase in M1 and M2 FC to a level higher than at rest (Figure 4D and Supplementary Figure 4Div & v). Therefore, cerebral cortical activity and FC differ prior to and after movement between motorized and spontaneous locomotion, often featuring differential involvement of anterior M2 nodes.

### Effects of speed and other behavioral parameters

In the above analyses, we used PLSR to remove the effects of motorized treadmill or freely-moving disk speed and other behavior parameters (see Methods), before comparing behavior periods or experimental conditions. The results of these initial regressions are isolated from the effects of each of these parameters on activation and FC (Figure 5 and Supplementary Figure 6). Therefore, we also investigated the effects of different locomotion speeds on cortical engagement during steady-state walk in each condition. During motorized and spontaneous steady-state walk, fluorescence decreases globally with increasing speed (Figure 5A and B). While the decreases in FC are uniform for motorized walk, spontaneous walk shows larger decreases in lateral M1 and S1 FC (Figure 5B). In both conditions during steady state walking, M2 activation and FC are similar in levels to other cortical regions. Acceleration rate was available in only the spontaneous walk period and had no significant effect on either activation or FC (data not shown). We also investigated the effect of increasing pupil diameter on all behavior periods, a commonly used measure of arousal^10^, and found it has a globally decreasing effect on activation and FC in most periods in both conditions (Supplementary Figure 6). The results further emphasize that the differences in anterior M2 FC between spontaneous and motorized locomotion occur primarily in the transition state and do not occur with changes in locomotion parameters during steady-state walk.

**Figure 5.**
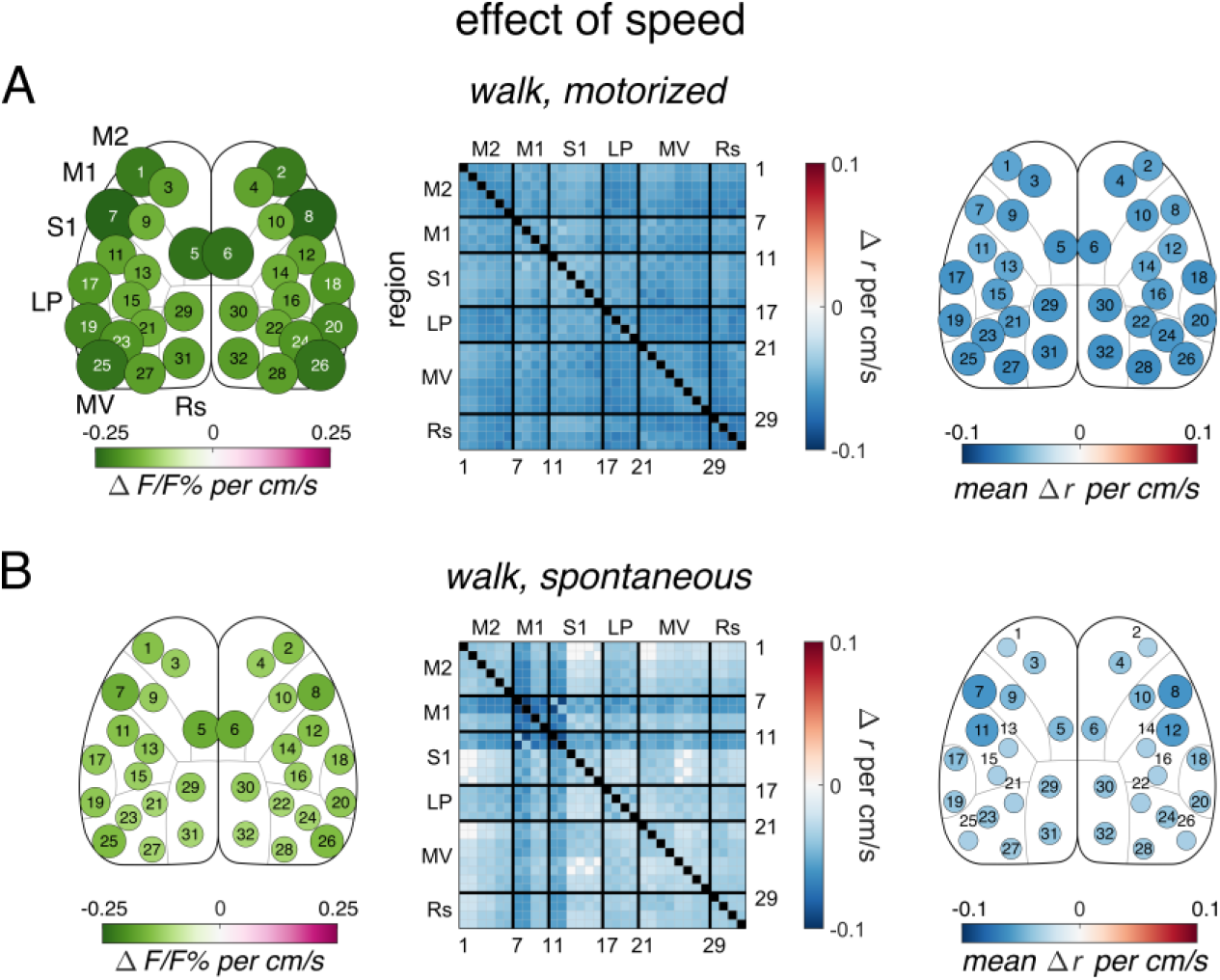
Effects of locomotion speed on cortical connectivity. **A**. Effect of speed on fluorescence and FC in motorized steady-state walk calculated with PLSR. **B.** Similar to A, but for spontaneous locomotion steady-state walk.

### Accelerating and decelerating periods

We were also interested in cortical engagement during locomotion transition states beyond simply starting and stopping. The motorized task includes accelerating and decelerating periods in which the mouse changes from one steady-state walking speed to another without first returning to rest (see Methods). Because the effects of various behavior parameters on the cortical FC are removed via regression, these periods can be interpreted as locomotion transition states without the influence of confounding variables, such as speed or acceleration rate. Unfortunately, these distinct transitions are not available in the spontaneous condition, although the most analogous measure, the effect of acceleration rate during steady-state walk, showed no significant effects on cortical activation or FC, as discussed above (data not shown). However, it is still useful to observe the effects of the accelerating and decelerating periods in the motorized condition to better understand cortical dynamics during these transitions in externally-driven locomotion.

The patterns of cortical activation and connectivity in accelerating and decelerating transitions are not only different from each other but also dissimilar from those in starting and stopping (Figure 6). Cortical activation decreases from steady-state walk levels when the motorized treadmill accelerates or decelerates (Figure 6Ai and iv). Activation remains depressed until after a new steady-state walking speed is reached (post-accelerating or post-decelerating periods; Figure 6Aii and v), before returning to steady-state levels (Figure 6Aiii and vi). However, FC differs between the accelerating and decelerating conditions. While both accelerating and decelerating are characterized by general decreases in FC (Figure 6Bi and iv), the largest decreases occur in M2 nodes in the decelerating period (Figure 6Biv). Then, when the new steady-state is reached, the network shows opposite responses between the post-accelerating or post-decelerating period, which are most prominent in FC between and among M1 and M2 nodes. In post-accelerating (a faster speed is reached) FC decreases (Figure 6Bii), while it increases in post-decelerating (a slower speed is reached; Figure 6Bv). The network then returns to the walk state (Figure 6Biii & vi). Acceleration rate has a global positive effect on the activation of the accelerating period, but there is no significant effect on the activation of the decelerating period or on FC in either (data not shown). Direct comparisons between the accelerating and decelerating periods (Figure 6C) confirm that M2 FC is higher in the accelerating period, indicating M2 FC decreases in only the decelerating period compared to walk. Also, activation drops further in the decelerating period in most nodes, but especially in M2 nodes (nodes 1-6) and posterior-lateral nodes (nodes 19-20 and 25-26; Figure 6C). These results show that accelerating and decelerating are different transition states from starting and stopping and that M2 has a role in the control of a wide range of locomotion transition types.

**Figure 6.**
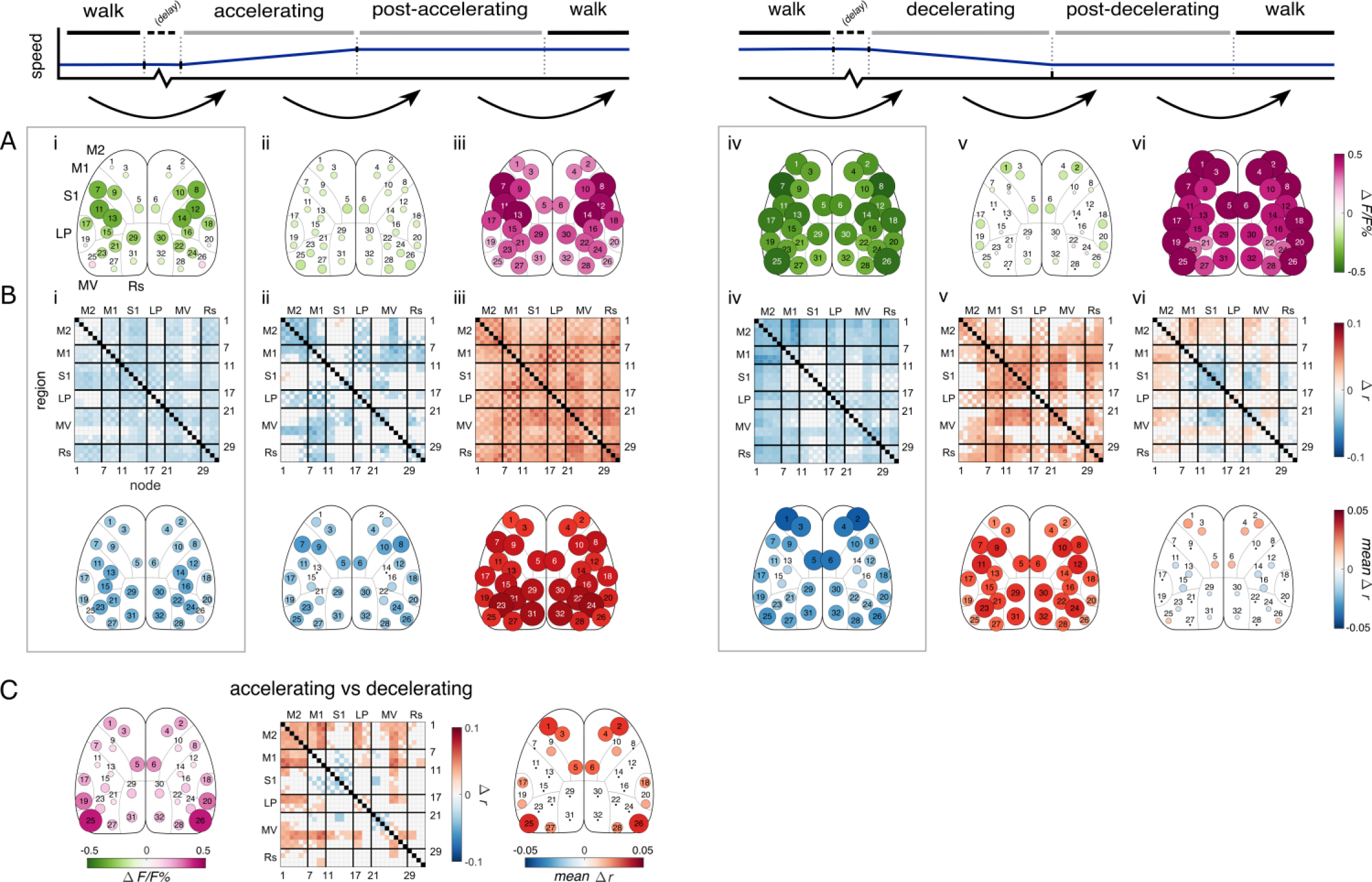
Accelerating and decelerating periods. **A.** Significant change in activation across sequential periods of motorized treadmill accelerating and decelerating periods, calculated and displayed as in Figures 2 & 3.Black arrows indicate the change in behavior periods being compared (i-vi). Positive values indicate increases in fluorescence or correlation across the change in behavior period. Delay periods are not included. Dotted boxes around accelerating (i) and decelerating compared to walk (iv) drawn for emphasis. **B.** Similar to A, but for FC. **C.** The significant difference in fluorescence (left) and correlation (middle and right) between the accelerating and decelerating periods (accelerating minus decelerating). Positive values indicate higher fluorescence or functional connectivity in the accelerating period.

## Discussion

We used PLSR to evaluate the evolution of cortical activation and FC networks through stages (called “periods”) of both motorized and spontaneous locomotion, and to evaluate the effects of individual behavior parameters on the cortex within each period. To the best of our knowledge, neither the activity nor the FC of the mouse dorsal cerebral cortex as a whole have been observed during externally-driven locomotion. Five major findings resulted from this study: First, we confirmed previous results in spontaneous locomotion that a distinct locomotion correlation pattern emerges, beginning at locomotion onset, that persists through steady-state walk until locomotion offset ^6^. This pattern is defined by increases in M2 FC and decreases in FC of all other regions. Second, externally-driven, motorized locomotion shows lower M2 FC during starting, stopping, and steady-state walk than internally-driven, spontaneous locomotion. However, motorized locomotion is characterized by increased activation during starting and decreased activation during stopping. Third, the preparatory periods before the onset of motorized and spontaneous locomotion result in very different activation and correlation patterns. The preparatory period before motorized starting shows high activation across the cortex with little change in FC, while the spontaneous preparatory period shows decreases in M2 FC but little change in activation. Instead, in spontaneous pre-start there is widespread ramp-up in activation and FC of all other regions until spontaneous locomotion onset, with no ramp in motorized pre-start. Fourth, activation and FC globally decrease with increasing locomotion speed in both the motorized and spontaneous conditions, but spontaneous locomotion shows additional modulation in M1. Finally, transitions in speed show differential effects depending on if speed is increasing or decreasing. Decreases in M2 FC occurs in the decelerating period, but not the accelerating period. In addition, the spatial patterns of activation and FC are not correlated, providing two modes of encoding and controlling cortical neural dynamics. Together, these results show externally-driven locomotion utilizes less cross-cortical integration than internally-driven, with M2 playing a differentiating role in the control of internally- and externally-driven transitions.

### Cortical activation in motorized and spontaneous locomotion

Both motorized and spontaneous locomotion exhibit widespread increases in fluorescence at the start of locomotion (Figure 2, Supplementary Figure 1). This activation is substantially larger in the motorized condition than in the spontaneous (Figure 3Aii) and persists into the post-starting period (Figure 2Aii, Supplementary Figure 4Aii). In both conditions, this initial activation eventually decreases over time as the mouse enters steady-state walk, similar to previous reports ^6, 10, 11^. This greater activation in motorized locomotion is striking and may be due to the increased computations required by the cortex to process incoming sensory information and develop an appropriate motor response. These computations begin before locomotion onset (Figure 4Ai) and accordingly diminish to equal those of spontaneous once steady-state walk is reached (Figure 2Aiii). In addition, motorized stopping leads to a decrease in activation below levels seen at rest (Figure 2Aiv), even as activation in spontaneous stopping remains elevated (Figure 2Ciii). This is a surprising finding, possibly due to an overall inhibition of cortico-cortical information exchange and a higher dependence on subcortical circuitry as slowing and stopping occur. Irrespective of the mechanism, these marked differences in activation emphasize that internally- and externally-driven locomotion utilize the dorsal cortical circuity quite differently.

### General decrease in functional connectivity in motorized and spontaneous locomotion

Here, we confirm our previous findings ^6^ that locomotion exhibits a widespread decrease in cortical FC compared to rest, even though neural activity in these areas increases ^12–15^. The exception to this global FC decrease is the increase in correlation of M2 nodes with nodes across the cortex, beginning at locomotion onset and persisting until locomotion offset. This occurs in both spontaneous and motorized locomotion and strongly supports our previous results ^6^.

Each of the cortical regions imaged show task-specific activity during walking, such as adjustment of gait in M1 ^16–19^, encoding of obstacles to be avoided in parietal cortex ^20–23^, and modulation of sensory encoding in somatosensory and visual cortices ^24–27^. Therefore, each of these regions engage in unique roles, and the proportion of *shared* fluorescence activity across them likely decreases. These individual roles may underlie the global decrease in correlations between cortical regions observed in both locomotion conditions.

### M2 in internally- and externally-driven locomotion

In our experiments, M2 FC differs between the internally- versus externally-driven conditions during locomotion transition. During starting, M2 FC is significantly higher in the spontaneous over the motorized case (Figure 3Bi). There is evidence suggesting locomotion-dependent modulation across the dorsal cortex originates from M2 ^15, 28–34^. Therefore, the increase in M2 FC observed at starting in both conditions may reflect a movement signal from M2 to other cortical regions. While this widespread modulation by M2 has not been investigated in externally-driven locomotion in rodents, activity in homologous cortical areas in primates have been implicated in the control of internally-driven movement more so than in externally-driven movement ^1–3, 35^. Therefore, it follows that cortical modulatory signals from M2 would be reduced at the onset of externally-driven locomotion, as seen here.

In stopping, M2 FC is again higher in the spontaneous condition than in motorized (Figure 3Biv). M2 FC increases compared to steady-state walk in spontaneous locomotion, but decreases in motorized (Supplementary Figure 4Biv and Diii). This is mirrored by M2 activation (Supplementary Figure 4Aiv and Ciii). Furthermore, after spontaneous locomotion offset only, M2 activation remains elevated (Figure 4D). Prefrontal cortical or M2 activity is thought to act as a “stop” control signal to subcortical structures to slow or inhibit locomotion ^36–38^. Our results support such a role for M2 in locomotion inhibition in spontaneous locomotion, marked by increases in M2 activation and FC, and this role is reduced in the externally-driven motorized treadmill case.

Notably, freezing-of-gait (FOG), a debilitating symptom of severe Parkinson’s disease, is partially linked to dysregulation of circuits involving the supplementary motor area (SMA) ^39–45^. Moreover, FOG is most likely to occur during gait transitions, not steady-state locomotion, and freezing during these transitions can be alleviated by guidance with an external cue ^46, 47^. This knowledge combined with our results supports the hypothesis that suggests upstream motor cortices such as SMA are fundamental to the differential control of internally- and externally-driven locomotion transitions.

### Locomotion preparation in internally- and externally-driven locomotion

In motorized locomotion, in addition to the widespread increase in activation discussed above, there is a decrease in M1 FC in the pre-start period (Figure 4A). Preparatory activity in M1 between warning and “go” cues is well documented ^48, 49^. Spontaneous locomotion, in contrast shows a decrease in M2 FC in the pre-start period, most prominently in the most anterior nodes (Figure 4B). This anticipatory change in FC may reflect motor planning ^2, 50, 51^ and locomotion initiation ^40, 41, 44, 45^ processes associated with upstream regions. As observed here, changes in M2 function can occur up to several seconds before movement onset ^2, 34, 51, 52^. M2 FC switches from a decreased to increased state at locomotion onset, perhaps reflecting a change from isolated motor planning to the delivery of motor information from M2 to the rest of the cortex.

Additionally, a large increase in activation and correlation occurs in all but anterior M2 nodes as locomotion onset approaches only in the spontaneous locomotion condition. The increase in FC does not depend on the increase in activity^6^. This pattern resembles the “ramp-up” summation pattern of increasing neural activation found in primate frontal and parietal regions in primates and across the cortex in rodents before decision-based, self-timed movements ^10, 49, 52–56^. Therefore, the observed ramp-up in correlations may reflect similar decision-making neural circuitry before spontaneous locomotion.

### Different effects of speed and other behavior parameters between internally- and externally-driven locomotion

The speed of locomotion has primarily global effects on both activation and correlation in steady-state walk; as speed increases, correlations decrease between most node pairs, in both motorized and spontaneous locomotion. This disagrees with previous findings that locomotion-dependent activity and modulation in a variety of cortical regions scales positively with running velocity ^6, 9, 44, 57, 58^. However, other studies have reported negative or no correlation between pyramidal cell activity and locomotion speed ^10, 11, 59^.Variability in reports may be due to the class or depth of the neurons recorded ^59^ or the behavior periods analyzed ^10, 11^. Here, we record activity primarily from excitatory pyramidal neurons of layer II/III (see Methods) while many of these previous studies record from layer V, which may result in distinct activation patterns ^60, 61^. Furthermore, we are unaware of any other studies that separate the contributions of speed, acceleration, and arousal to activation or FC as done here, which may mask a negative correlation.

Spontaneous locomotion also induces additional decreases in correlations with speed in lateral M1 and somatosensory (S1) nodes that are strongly associated with limb movement ^62, 63^. Both M1 and S1 activity changes with the phase of the step cycle in humans ^64–66^ and in cats ^17, 18, 67^. It is possible that the FC of these nodes does not change as much with speed in the motorized condition because externally-driven, simple steady-state walk requires little cortical input for gait adjustment ^68^. Additional experiments using equivalent speeds are required to clarify this difference in the cortical network.

### Accelerating and decelerating transition periods

During the motorized treadmill condition we introduced controlled changes in speed, the accelerating and decelerating periods. Using PLSR, we removed the effects of the beginning and end speeds, acceleration rate, and arousal from the correlation network in these periods, leaving only the effects of the transition state itself. Unexpectedly, activation decreases when accelerating or decelerating from steady-state walking. Previous electrophysiology studies during changes in gait pattern suggest cortical activation would increase during these periods ^69–71^. However, these studies record from pyramidal neurons in the deep layers of the cortex, specifically layer V, which project to subcortical structures. Here, our recordings predominantly detect activation of pyramidal neurons in layer II/III (see Methods), which project across cortical regions. These populations show distinct activation patterns ^60, 61^. Therefore, our results support that cortico-cortical activity decreases during accelerating or decelerating periods, even as subcortical-projecting neural activity may increase.

Interestingly, the accelerating and decelerating periods have different effects on FC patterns. The decelerating period shows decreases in M2 correlation pairs that are not present in the accelerating period, especially between other M2 nodes and M1 nodes, suggesting decelerating uses a distinct neural mechanism than accelerating. We see similar decreases in M2 correlations during motorized treadmill stopping (Supplementary Figure 4Civ). Human subjects walking on a treadmill find it easier to decrease their stepping rates over increasing them in response to an external cue ^38, 72^, and this difference in effort is linked to prefrontal cortex activity that may act as a locomotion “stop” mechanism ^37, 38^. With this evidence taken into account, our findings support that M2 is involved in a locomotion inhibition mechanism utilized in decelerating but not in accelerating under similar conditions.

### Lack of changes in retrosplenial cortex connectivity

Notably, these results do not reproduce the increases in the FC between retrosplenial nodes and the rest of the cortex during locomotion found in previous experiments ^6^. Here we observe significant increases only between retrosplenial and visual nodes during starting in both motorized and spontaneous locomotion, which falls during steady-state walk to levels similar to those at rest. Another group has also reported increases in correlation between retrosplenial and visual cortex during locomotion, although they found these increases persist through steady-state walk ^12^. The discrepancies between our previous and current findings likely originate from the differences in the processing and analytical methods implemented. Importantly, here we used a more complete hemodynamic correction and blood vessel removal procedure and regressed out the effects of various behavior parameters before comparing correlations across periods. Additional experiments are necessary to further clarify the changes in retrosplenial cortex FC in locomotion.

### Limitations

There are limitations to the study presented. Importantly, we acknowledge that locomotion during head-fixation is not identical to naturally occurring, free-ranging locomotion. We strove to minimize the effects of head-fixing by habituating the animals to the behavioral setup and training them to walk comfortably on the motorized treadmill over time at least 2 weeks before beginning experiments (see Methods). Additionally, both the motorized and spontaneous locomotion conditions were performed with head-fixation, ensuring differences between the two conditions are due to differences in locomotion generation, and not because of differences between head-fixation or free motion. Unfortunately, head-fixation reduced the upper limit of the speed of the motorized treadmill that the mice could safely walk. Thus, the locomotion speeds in the motorized condition were lower than the average speed when mice walked spontaneously. We corrected for this discrepancy by first removing the effect of locomotion speed on the cortical correlations before any comparisons between conditions were calculated. Therefore, the effects of speed differences on correlation across conditions were limited.

Generally, M2 in rodents is accepted to include functionality homologous to premotor cortex and SMA in primates ^33, 73, 74^. Our results pertaining to M2 are most prominent in the anterior nodes, nodes 1 and 2. Much of the research to date involving M2-like regions in locomotion, or in the differences between internally- and externally-driven movement, has found similar effects in premotor cortex and SMA, as described above. Therefore, we do not attempt to link our findings to any particular homologous region outside of rodents.

### Conclusion

In conclusion, cortical FC and activation are markedly different between externally- and internally-driven locomotion transitions. Anterior M2 FC is heightened in internally-driven transitions compared to externally-driven, suggesting this region plays an important role in the cortical-level control of these two behavioral states.

## Methods

### Animals and surgical setup

Seven Thyl-GCaMP6f mice were used in these experiments (3 female, 4 male; The Jackson Laboratory). Thy1-GCaMP6f mouse lines express GCaMP6f ^75^ in layers II/III and V cortical excitatory pyramidal neurons ^76^, allowing for wide-field Ca^2+^ fluorescence imaging of predominantly layer II/III excitatory neurons ^77^. Mice were surgically implanted with See-Shell polymer windows over the dorsal cerebral cortex ^78^ as described previously ^6, 79, 80^. Mice were housed in a 12hr-12hr light-dark cycle, with times of darkness (the period of mouse wakefulness) occurring during the day when experiments were performed. All animal studies were approved by and conducted in conformity with the Institutional Animal Care and Use Committee of the University of Minnesota.

### Behavioral task

Mice were trained to walk on a motorized disk treadmill that introduced controlled locomotion speeds, onsets, offsets, and changes in speed (Figure 1C, top). The treadmill was powered by a stepper motor (MotionKing, Nema 17 2-phase stepper motor) driven by a high-current stepper motor driver (Texas Instruments, DRV8825 Stepper Motor Controller), and controlled with an Arduino Due microcontroller (Arduino) with custom Arduino code (Arduino IDE v1.8).

The behavior task was composed of 5-minute trials divided into stages of randomly varying length in time (3 to 25 s; Figure 1C, top). Each trial began and ended with the motor at rest (speed = 0 cm/s). Intermediary stages ran the treadmill at one of 5 randomly assigned speeds: 0 (rest), 2.25, 2.77, 3.33, or 3.88 cm/s. At the beginning of each stage, the treadmill transitioned to the new speed at one of two possible randomly selected acceleration rates: 0.56 or 1.11 cm/s/s for starts and stops, and 0.28 or 1.11 cm/s/s for accelerating or decelerating periods. The speeds and accelerations were chosen by observing the comfortable walking speed of head-fixed mice, which are slower than mice comfortably attain in spontaneous locomotion (see West et al., 2022).

Speeds and acceleration rates also accounted for the hardware parameters of the stepper motor so that stages began and ended on 1.0- or 0.5-second intervals. Transition periods were labeled for analysis based on the beginning and ending speeds of the transition (starting: 0 cm/s to any other speed; stopping: any non-zero speed to 0 cm/s; accelerating: any non-zero speed to a different, faster speed; decelerating: any non-zero speed to a different, non-zero, slower speed; maintaining: when both the beginning and ending speeds are the same and no transition in speed occurs). Auditory warning cues were presented to the mice before the treadmill changed speed to reduce surprise or startle and the associated brain activity. After the mouse completed a training paradigm that slowly increased the latency between the warning cues and the onset of a transition, warning cues were presented 5 s before the transition and lasted a total of 1 s. Different tone patterns based on mouse auditory ranges ^81^ indicated which of the 5 possible transitions would occur (Supplementary Table 4) and were played simultaneously on two piezoelectric buzzers (TDK, PS1240P02CT3). Playing these high-frequency tones while simultaneously controlling the stepper motor at a smooth rate required the processing rate of the Arduino Due (84 MHz). The periods between the tones and the onset of the transition were analyzed as “delay periods.” The 1 s in which the tones actively sound and the 1 s immediately after the tones were thrown out during analysis to remove activity resulting from direct auditory processing of the tones, leaving the 3 s immediately before the onset of the transitions.

After a minimum of 7 days recovery post-surgery, mice were habituated to the behavioral setup (Figure 1A) over at least 4 days, as described previously ^6, 80^. After habituation, mice were trained over at least 10 days by gradually increasing the locomotion speed and complexity of the motorized treadmill task. Each mouse underwent 10 trials of the motorized task during Ca^2+^ fluorescence imaging per day, for a total of 50 minutes per day. In addition, mice were recorded for 5 trials of spontaneous locomotion for a total of 25 minutes per day, immediately before or after the motorized task. More than 90 hours of locomotion imaging was collected, resulting in ∼1500 instances of each transition type (Table 1). One mouse (mouse #4) only rarely walked spontaneously (Table 1) and was not included in the spontaneous locomotion analysis. Behavior in both conditions was reported and recorded via the Arduino serial monitor and PuTTY terminal software.

**Table 1.**
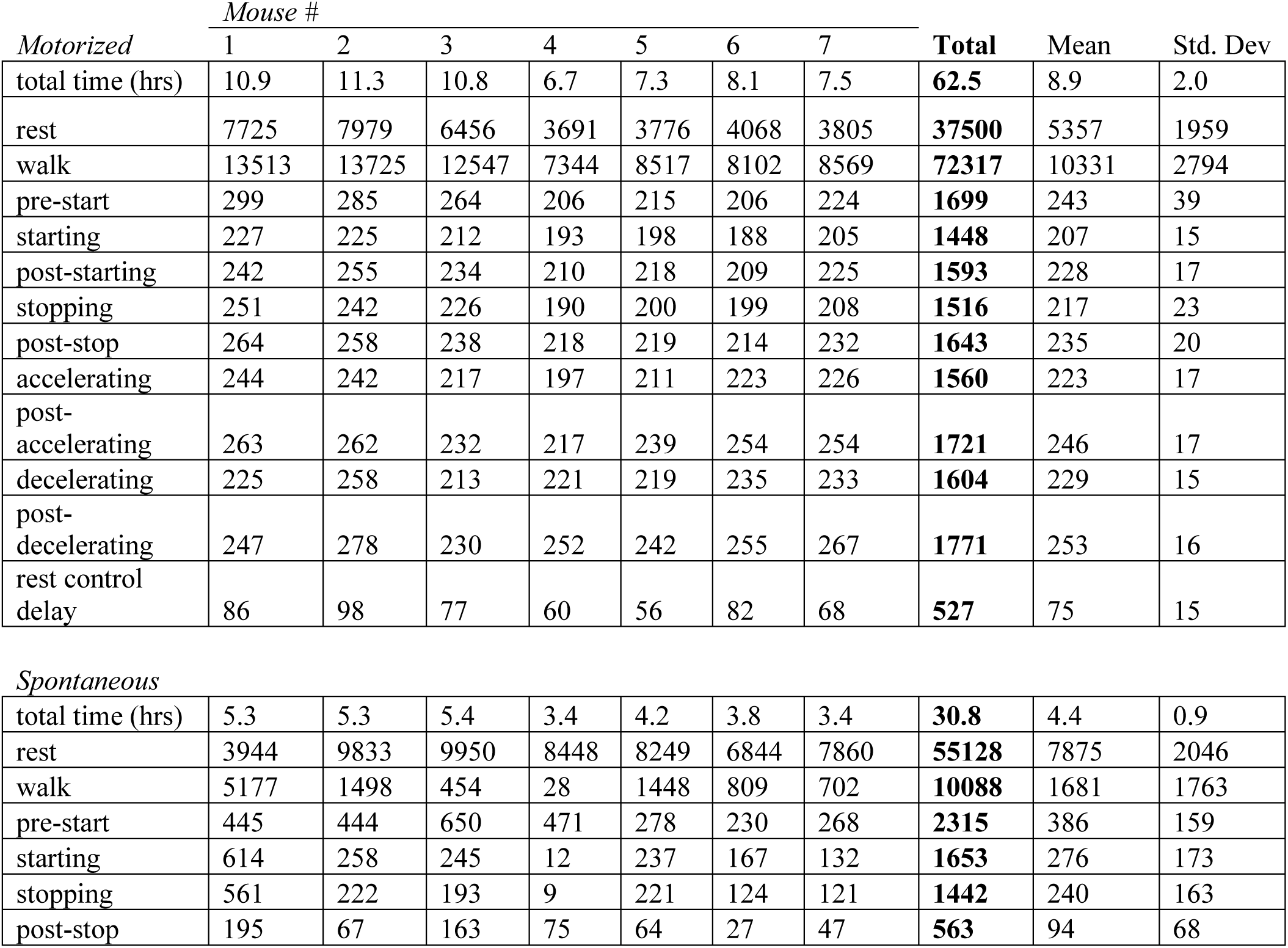
Instances per behavior period.

| *Motorized* | *Mouse #* |  |  |  |  |  |  | **Total** | Mean | Std. Dev |
| --- | --- | --- | --- | --- | --- | --- | --- | --- | --- | --- |
|  | 1 | 2 | 3 | 4 | 5 | 6 | 7 |  |  |  |
| total time (hrs) | 10.9 | 11.3 | 10.8 | 6.7 | 7.3 | 8.1 | 7.5 | **62.5** | 8.9 | 2.0 |
| rest | 7725 | 7979 | 6456 | 3691 | 3776 | 4068 | 3805 | **37500** | 5357 | 1959 |
| walk | 13513 | 13725 | 12547 | 7344 | 8517 | 8102 | 8569 | **72317** | 10331 | 2794 |
| pre-start | 299 | 285 | 264 | 206 | 215 | 206 | 224 | **1699** | 243 | 39 |
| starting | 227 | 225 | 212 | 193 | 198 | 188 | 205 | **1448** | 207 | 15 |
| post-starting | 242 | 255 | 234 | 210 | 218 | 209 | 225 | **1593** | 228 | 17 |
| stopping | 251 | 242 | 226 | 190 | 200 | 199 | 208 | **1516** | 217 | 23 |
| post-stop | 264 | 258 | 238 | 218 | 219 | 214 | 232 | **1643** | 235 | 20 |
| accelerating | 244 | 242 | 217 | 197 | 211 | 223 | 226 | **1560** | 223 | 17 |
| post-accelerating | 263 | 262 | 232 | 217 | 239 | 254 | 254 | **1721** | 246 | 17 |
| decelerating | 225 | 258 | 213 | 221 | 219 | 235 | 233 | **1604** | 229 | 15 |
| post-decelerating | 247 | 278 | 230 | 252 | 242 | 255 | 267 | **1771** | 253 | 16 |
| rest control delay | 86 | 98 | 77 | 60 | 56 | 82 | 68 | **527** | 75 | 15 |
| *Spontaneous* |  |  |  |  |  |  |  |  |  |  |
| total time (hrs) | 5.3 | 5.3 | 5.4 | 3.4 | 4.2 | 3.8 | 3.4 | **30.8** | 4.4 | 0.9 |
| rest | 3944 | 9833 | 9950 | 8448 | 8249 | 6844 | 7860 | **55128** | 7875 | 2046 |
| walk | 5177 | 1498 | 454 | 28 | 1448 | 809 | 702 | **10088** | 1681 | 1763 |
| pre-start | 445 | 444 | 650 | 471 | 278 | 230 | 268 | **2315** | 386 | 159 |
| starting | 614 | 258 | 245 | 12 | 237 | 167 | 132 | **1653** | 276 | 173 |
| stopping | 561 | 222 | 193 | 9 | 221 | 124 | 121 | **1442** | 240 | 163 |
| post-stop | 195 | 67 | 163 | 75 | 64 | 27 | 47 | **563** | 94 | 68 |

### Fluorescence imaging

All fluorescence imaging was performed with dual-wavelength illumination to minimize the effects of hemodynamics on the Ca^2+^ signal ^6, 28, 79, 80^. Briefly, mice were head-fixed beneath the fluorescence microscope on the motorized treadmill or freely moving disk (Figure 1A) for imaging of the dorsal cerebral cortex (Figure 1B). Ca^2+^-dependent (470 nm, blue light) and Ca^2+^-independent (405 nm, violet light) GCaMP6f signals were captured on consecutive frames at 40 frames per second (fps; 20 fps per channel). Imaging was performed in 5-minute image stacks (1 per behavior trial), each 6000 frames in length per color channel. Behavior was captured with infrared (IR) cameras ^6^, recording full body movement at 40 fps and the eye at 20 fps. Individual frames of the fluorescence imaging, the IR behavior cameras, and the onset of the Arduino-controlled behavior task were precisely timed by a custom program in Spike2 (Cambridge Electronic Design Limited, version 5.21).

### Fluorescence preprocessing

All preprocessing was performed using custom code in MATLAB. For each recording day, the first blue-channel image of the first image stack was chosen as a representative image for that recording session. For each mouse, the clearest and most centered image of all recording sessions was chosen as that mouse’s overall representative image. This image served as the reference image for registration and drawing brain masks (see below). The Spike2 program checked that the first image in a stack was from the blue channel (i.e., all odd images were blue channel). All blue channel images were spatially registered to that recording session’s representative image using the *dftregistration* MATLAB function ^82^. The resulting transformation matrices for each blue image were applied to the corresponding violet images. Next, images were registered across recording sessions within mice using MATLAB Image Processing Toolbox function *imregtform* using an affine transformation. Brain masks were drawn for each mouse so only the pixels corresponding to brain were kept for further analysis.

A 7 Hz low-pass temporal filter (5^th^ -order Butterworth) was applied with the MATLAB Signal Processing Toolbox function *filtfilt*. The effects of hemodynamics and other non-neuronal sources were removed from the fluorescence signal using the violet-channel images. Within each stack, the activity of each pixel in the blue channel was regressed against the activity of the corresponding pixel in the violet channel ^28^, using a simple linear regression with intercepts via the MATLAB built-in function *regress.* The residuals of these regressions were kept as hemodynamic-corrected images, which were used in all subsequent steps of analysis.

### Fluorescence spatial segmentation

Spatial segmentation of the dorsal cerebral cortex was performed with singular value decomposition (SVD) compression and independent component analysis (ICA) as described in West et al. (2022). Several changes were made, however, due to the enhanced quality of the imaging resulting from modifications to the fluorescence preprocessing. All fluorescence imaging per mouse was compressed to 500 SVD components using a randomized algorithm ^83^. The Joint Approximation Diagonalization of Eigenmatrices (JADE) algorithm for ICA was used to find 100 independent components (ICs) Resulting ICs were thresholded to spatial domains of 150-5000 pixels in size. Each IC was thresholded so that pixels with z-scored weights below 3.5 were set to equal 0, and ICs of at least 150 contiguous remaining pixels were kept. If no domains of 150 pixels survived, this initial z-score threshold was reduced to 2.5, and any surviving domains of 150 pixels were kept. If initial thresholding instead generated domains greater than 5000 contiguous pixels in size, which occurred rarely, a second z-scoring was applied to the remaining data in the domain and a threshold of 1.0 was applied. After thresholding was complete in all cases, if more than one domain of at least 150 contiguous pixels occurred in a single IC, those domains were separated and treated as different in the following analysis steps. Resulting domains were visually inspected, and any domains corresponding to artifacts such as blood vessels were discarded. The dual-wavelength hemodynamic correction procedure may not remove all artifacts in the fluorescence signal due to the fluctuation of blood vessel diameter, particularly during the physical exertion of locomotion. To avoid such signal contamination in the remaining IC domains, pixels that overlapped with blood vessels were manually removed.

IC domains were manually assigned node IDs according to domains that appeared consistently across mice (32 nodes were identified, see Figure 1D and E). Visual inspection revealed raw ICs that matched a given node ID but did not survive the thresholding process. The thresholding of ICs is admittedly somewhat arbitrary, focused on automated, time-efficient analysis, and this occasionally discarded an IC in a given mouse that, on visual inspection, corresponded to a node present in all mice. Excluding nodes in individuals could detrimentally affect conclusions, therefore these ICs were manually cropped, thresholded, and included in further analysis.

### Behavior data extraction and preprocessing

Motorized treadmill task parameters from the Arduino microcontroller output were extracted and analyzed using custom MATLAB code. The entirety of the motorized trials was subdivided and sorted into the possible behavior periods (see Figure 1C) and labeled with the speed of the treadmill during that period. Transition periods were labeled with both the speeds at the beginning and end of the transition as well as the transition acceleration rate. Disk speed in the spontaneous locomotion trials was extracted, processed, and divided into behavior periods as described in West et al. (2022). Disk acceleration was also calculated.

To extract pupil diameter, the outer edge of the pupil was tracked using DeepLabCut ^84^. Eight equidistant points were labeled and tracked around the pupil edge. For each frame, a circle was fit through all points using the Pratt method ^85^, and the diameter of that circle was calculated and normalized to the mouse’s maximum pupil diameter for that recording day. The pupil diameter was not always detectable during recordings, due to eye blinks/closing or drift in camera position. As partial least squares regression (PLSR, see below) cannot directly incorporate missing values, trimmed score regression (TSR) was used to estimate the missing pupil diameters using the available behavior parameters (speed, acceleration rate, and duration) and fluorescence data ^86^. This method preserves the correlation structure of the data ^86–88^ and is described in detail below.

### Fluorescence timeseries extraction, sliding windows, and correlations

To obtain fluorescence timeseries for each node, the mask of the corresponding IC for a given mouse was applied to the preprocessed fluorescence images. The IC weights within the mask were used to create a weighted mean fluorescence value per frame. All resulting fluorescence timeseries were converted to the percent change in fluorescence divided by baseline fluorescence (ΔF/F%), in which the baseline is the mean fluorescence across all recording sessions for each mouse and node. Timeseries were then segmented and grouped by behavior period within each mouse.

To investigate the effect of the amount of time since the mouse had begun a given period (i.e., since having begun a speed transition, reached a new speed, or been presented with a warning cue), fluorescence timeseries were subdivided using a sliding window with a 1.0 s (20 frame) window size and 0.25 s (5 frame) step size and an “instantaneous” duration value equal to the time in seconds at the center of each window. Pearson correlation coefficients were then calculated on each 1.0 s window between the timeseries of all possible node pairs, creating correlation matrices for each window (20 time points in each calculation, 496 total correlation pairs). The Fisher z-transformation was applied to all coefficients to create approximately normally distributed datasets ^89, 90^. This analysis did not investigate differences between activity of the left and right hemispheres, so the correlations of anatomically homologous nodes on the left and right hemispheres (nodes falling at the same anatomical location on the contralateral hemisphere) were averaged. Ipsilateral and contralateral relationships were preserved, however, by only averaging equivalent ipsilateral or contralateral correlations.

The sliding windows were also applied to the disk or treadmill speed and acceleration as well as to the eye pupil diameter over time. For all analyses, the acceleration rate was converted to the absolute value of the recorded treadmill or disk acceleration rate. These parameters were averaged within 1.0 s time window (20 time points) to create a single parameter value per corresponding fluorescence correlation matrix.

### Partial Least Squares Regression (PLSR)

Partial Least Squares regression (PLSR) was used to relate locomotion behavior to the FC of the dorsal cerebral cortex. PLSR initially utilizes dimensionality reduction to optimize the signal-to-noise ratio of the relationship between the independent and dependent variables before performing a regression ^91–93^. PLSR was performed using the MATLAB Statistics and Machine Learning Toolbox function *plsregress*^91^. Two levels of regressions were performed. The first were continuous regressions in which the behavior parameters (speed, acceleration rate, duration, and pupil diameter) were regressed against correlations between nodes within each behavior period. The second were categorical regressions in which the different behavior periods, coded as dummy variables, were regressed against the corresponding correlations between nodes. Correlations between nodes were the independent variable and behavior parameters or periods were the dependent variables.

The effects of behavior parameters on correlation between nodes within each behavior period were evaluated first. Each behavior parameter (speed, acceleration rate, duration, and pupil diameter) formed a dependent variable, and each possible correlation pair between nodes (496 correlation pairs) formed an independent variable. Each set of correlation pairs made up a single observation (i.e., the correlations between nodes from a single sliding window were an observation, with each subsequent sliding window forming a new observation). The behavior parameters were collected into the *n* x *p* dependent variable matrix *Y* (*n =* number of observations; *p* = number of behavior parameters, up to 4), and correlations were collected into the *n* x *m* independent variable matrix *X* (*n* = number of observations; *m* = number of correlation pairs, 496). The regression was performed as:

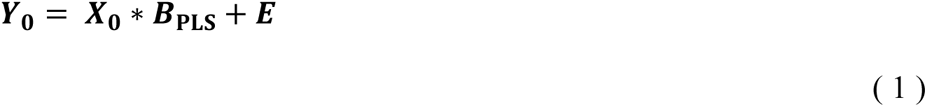

In which Y_0_ and X_0_ are Y and X normalized within each variable, respectively, *B*_PLS_ is the matrix of normalized regression slopes calculated by PLSR (*m* x *p*) and *E* is the matrix of residuals of the correlation data (*n* x *m*). Diagrams of the regression equations are provided (Supplementary Figure 2). A column of ones was included in *X_0_* to include intercepts in the regressions.

As described above, data for the pupil diameter was occasionally missing, and PLSR cannot directly incorporate data with missing values ^86^. Therefore, trimmed score regression (TSR) was used to estimate the missing pupil diameters for inclusion in PLSR, using the MATLAB function *plsmbtsr1* ^86^. The TSR was run with a number of principal component analysis (PCA) components that explained at least 85% of the variability of the combined matrix *M* = [*X_0_*, *Y_0_*], with a maximum of 10 components. TSR was run with a minimum convergence of 1 x 10^-10^ and 1000 maximum iterations. The estimated pupil diameter values were then incorporated into *X_0_* and *Y_0_*, and PLSR with the behavior parameters for each behavior period was performed.

To determine the number of PLSR components that provided the best prediction of *Y_0_* from *X_0_*, each regression was performed with 1 to 10 components, with 10-fold cross validation and 10 Monte Carlo repetitions. Cross validation was performed on subsets of observation that were contiguous in time. For each Monte Carlo repetition, the observations were listed in sequential order, then divided into 10 equal segments with the division points offset by a random amount. The cross-validated mean squared error (MSE) for the dependent variables was calculated for each possible number of PLSR components, and the Bayes Information Criterion (BIC) ^94^ was applied. The model that produced the lowest adjusted MSE was interpreted as the model that best explained the covariance between *Y_0_* from *X_0_*, and the number of components in the model was used in further analysis steps. The optimal number of components was found for each mouse and behavior period.

Regression of behavior parameters against correlations between nodes not only revealed the influence of those parameters on network FC, but also allowed the investigation of network FC across behavior periods without the variability in those parameters acting as confounding factors. The residuals of the behavior parameters regressions were kept for comparisons across behavior periods (Supplementary Figure 2C). Categorical regressions across behavior periods were performed between pairs of behavior periods of interest. In these regressions, Y was a *n* x 2 matrix of dummy variables (1, 0 or 0, 1) identifying each observation as belonging to one behavior period type or the other, again normalized by column.

Regressions were performed as above, with the additional step of stratifying the cross validation so that each train and test subset of the data contained a ratio of each behavior period equal to the original dataset.

Traditionally, *B*_PLS_ is calculated as regression coefficients that relate *Y_0_* to an additionally normalized version of *X_0_,* called *W*, and this is the *B*_PLS_ matrix calculated by the MATLAB *plsregress* function (see eq 1; de Jong, 1993). However, here we are interested in the expected change in the original matrix *X* as the parameters *Y* change. In addition to *B*_PLS,_ PLSR calculates the loadings of *X_0_* and of *Y_0_* onto the latent components, abbreviated here as *XL* and *YL*, respectively. *XL* is a *m* x *c* matrix (*m* = number of correlation pairs, *c* = number of PLSR components), while *YL* is a *p* x *c* matrix (*p* = number of behavior parameters, *c* = number of PLSR components). When PLSR optimization is included, expected change in *X* with *Y* is expressed as:

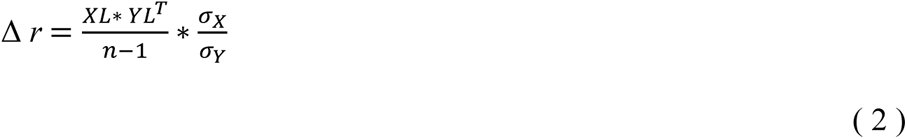

where σ_X_ and σ_Y_ are vectors containing the standard deviations of each variable in *X* and *Y*, respectively. This is equivalent to an PLSR-optimized matrix of regression slopes between the behavior variables and the correlations between nodes, here called the “change in correlation” matrix (Δ *r*). This gives the expected change of each node-node correlation pair, either as given behavior parameter changes within a behavior period per unit, or as one behavior period is compared to another. Changes in correlation matrices were averaged across mice. The resulting PLSR metrics are provided in Supplementary Tables 5 and 6. Generally, regressions against behavior parameters required fewer PLSR components and performed better than those across different behavior periods.

### PLSR on fluorescence timeseries

To determine if changes in FC could be linked to changes in neural activity, PLSR was also performed on the mean fluorescence time series from each node. These regressions were performed similarly to those performed on FC, as described above. Results from these regressions are reported as the change in fluorescence as a percent of the mean.

### Relationship between change in fluorescence and change in correlation

The relationship between the PLSR-calculated changes in fluorescence and FC was determined by calculating the correlation between these two measures. For each node, changes in fluorescence and FC were correlated across all the investigated categorical comparisons for each node. This was repeated for the behavior parameter comparisons (speed, acceleration rate, duration, and pupil diameter).

### Statistics

For each PLSR regression, null distributions were generated for each node within a mouse using a 1000-permutation test, then averaged across mice to create a single null distribution. P-values were then calculated with a two-tailed z-test using the MATLAB Statistics and Machine Learning Toolbox function *ztest* (α = 0.05) and adjusted for multiple comparisons with the False Discovery Rate (FDR) post-hoc correction method ^95^. Significance of the relationship between activation and FC was determined using a 2-tailed t-test with FDR correction with individual mice as replicates (degrees of freedom = 6, t-statistics shown in Table 2).

**Table 2.** T-statistics for activation-FC correlations (degrees of freedom = 6).

| **Nodes** | **Across periods** | **Speed** | **Acceleration rate** | **Duration** | **Pupil diameter** |
| --- | --- | --- | --- | --- | --- |
| **1-2** | 3.35 | 2.19 | -0.84 | -0.19 | 1.04 |
| **3-4** | 3.62 | 3.45 | -1.78 | 0.79 | 0.81 |
| **5-6** | 0.22 | 1.18 | -1.08 | 1.48 | 0.21 |
| **7-8** | -0.37 | 1.33 | -0.49 | 1.34 | 0.40 |
| **9-10** | 2.47 | 7.25 | -2.10 | 1.79 | 0.80 |
| **11-12** | -5.55 | 0.28 | -0.17 | 0.79 | 0.27 |
| **13-14** | 2.69 | 3.82 | -0.89 | 1.77 | 0.65 |
| **15-16** | -3.71 | 0.39 | 0.58 | 1.45 | 0.46 |
| **17-18** | 0.83 | 6.78 | -0.30 | 2.82 | 0.63 |
| **19-20** | -0.68 | 2.95 | 0.47 | 1.99 | 0.35 |
| **21-22** | 0.03 | 6.10 | 0.72 | 3.03 | 0.47 |
| **23-24** | 2.71 | 4.73 | -1.14 | 1.60 | 0.41 |
| **25-26** | 0.95 | 4.91 | 0.35 | 3.12 | 0.47 |
| **27-28** | 3.26 | 3.86 | -0.17 | 2.26 | 0.38 |
| **29-30** | 2.50 | 4.04 | 0.08 | 4.05 | 0.47 |
| **31-32** | 1.99 | 4.53 | -1.44 | 3.31 | 0.70 |

### Code availability

Code used in these analyses is available in the repositories listed in Table 3.

**Table 3.** Code repositories.

| **Use** | **Repositories** |
| --- | --- |
| Motorized treadmill task control | https://github.com/west0883/motorizedTreadmill_state |
| Spontaneous locomotion rotary encoder control | https://github.com/west0883/rotary_encoder_recording |
| Universal functions | https://github.com/west0883/code_common_to_pipelines |
| Motorized treadmill behavior segmentation | https://github.com/west0883/motorized_treadmill_behavior_pipeline |
| Spontaneous locomotion behavior segmentation | https://github.com/west0883/treadmill_encoder_pipeline_code |
| Fluorescence imaging preprocessing | https://github.com/west0883/preprocessing_pipeline_code |
| Fluorescence imaging spatial segmentation | https://github.com/west0883/MSI_SVD_compression_code<br>https://github.com/west0883/IC_pipeline_code |
| Fluorescence initial computations | https://github.com/west0883/fluorescence_analysis_pipeline |
| PLSR | https://github.com/west0883/partial_least_squares_pipeline_code |

## Supporting information

Supplementary Materials with Figures

## Data availability

Data generated and analyzed in this study are available from the authors upon request.

## Acknowledgements

We would like to thank Lijuan Zhuo for her assistance in animal surgeries. The Minnesota Supercomputing Institute provided high-processing computing, and the University of Minnesota University Imaging Centers (UIC, SCR_020997) provided 3D printing services of the cortical implants. This work was supported by National Institutes of Health (R01- NS111028 to T.J. E., P30- DA048742 to T.J.E., and T32-MH115688 to S.L.W., as well as by a grant from University of Minnesota Informatics Institute, which includes support from the University of Minnesota’s MnDRIVE Initiative (to S.L.W).

## Contributions of the authors

SLW developed the experimental design and hardware required, performed experiments, led the analyses and writing of the manuscript. MLG performed experiments and data collection. TJE was involved with the experimental design and writing of the manuscript as well as oversight of and secured funding for project.

