## Supplementary Materials with Figures for "Wide-field calcium imaging of cortical activation and functional connectivity in externally- and internally-driven locomotion"

### Supplementary Tables

**Table 4.** Warning tone parameters. Each tone consisted of 3 pitches in sequence, played for 310 ms with a 23 ms silent period in between to increase tone salience, for a total of 1.0 s. Different pitches were played to distinguish between the upcoming transitions.

| **Upcoming transition** | **Pitch in Hz** | | |
|---|---|---|---|
| | 1st pitch | 2nd pitch | 3rd pitch |
| starting | 10,000 | 10,000 | 10,000 |
| stopping | 4,000 | 4,000 | 4,000 |
| accelerating | 4,000 | (silence) | 10,000 |
| decelerating | 10,000 | (silence) | 4,000 |
| maintaining | 7,000 | (silence) | 7,000 |

**Table 5.** PLSR metrics

| **Metric** | **Regression type** | **Mean** | **Std. dev. across mice** | **Std. dev across regressions** |
|---|---|---|---|---|
| # PLSR components | *parameters* | **1.8** | 0.6 | 1.0 |
| | *periods* | **4.7** | 0.8 | 1.8 |
| Mean squared error | *parameters* | **0.95** | 0.02 | 0.05 |
| | *periods* | **0.38** | 0.14 | 0.27 |

**Table 6.** Network connectivity percent variance explained per behavior variable.

| **Behavior variable** | **Mean** | **Std. dev. across mice** | **Std. dev across regressions** |
|---|---|---|---|
| *Speed* | 9.5% | 2.1% | 6.3% |
| *Acceleration* | 6.4% | 1.7% | 2.8% |
| *Duration* | 8.2% | 1.7% | 7.7% |
| *Pupil diameter* | 9.8% | 4.4% | 2.2% |
| *Behavior periods* | 16.8% | 2.7% | 10.1% |

### Supplementary Figures

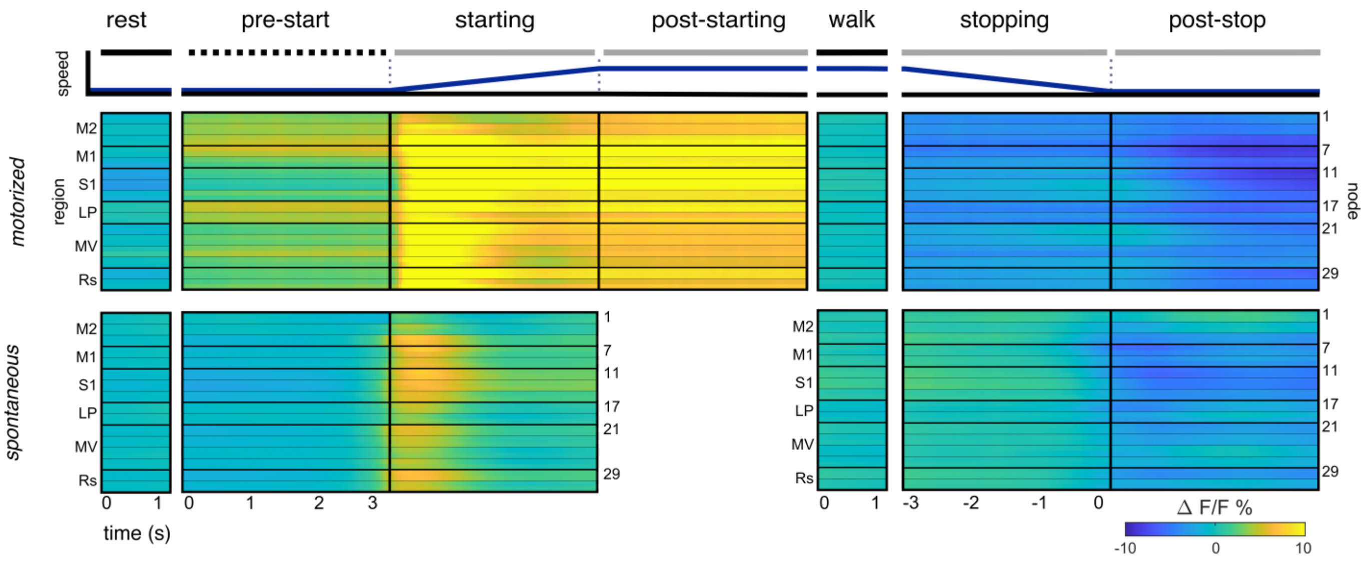


**Supplementary Figure 1. Activation in motorized treadmill and spontaneous locomotion**

ΔF/F% of nodes across locomotion behavior period for motorized treadmill (top) and spontaneous locomotion (bottom) conditions. There is no post-starting period defined in the spontaneous condition.

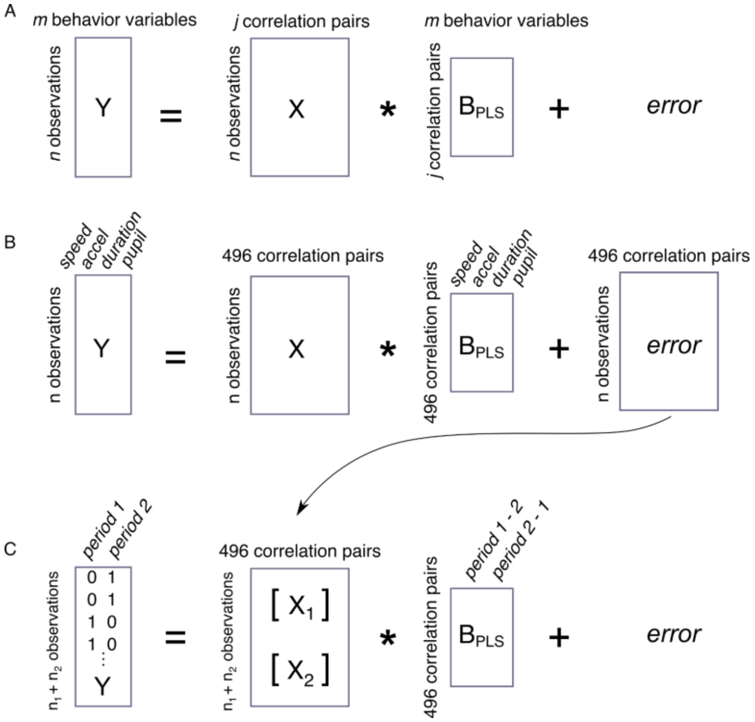


**Supplementary Figure 2. Diagram of partial least squares regression (PLSR) steps.**

**A.** Diagram of the general regression equation used. **B.** Diagram of the equation used for each behavior period regressing behavior parameters against FC. The error term (residuals) of these regressions is then carried forward as the node-node correlation data in the regressions across behavior periods (the X matrix in C), as indicated by the arrow. **C.** Regression equation used to calculate the change in FC across behavior periods.

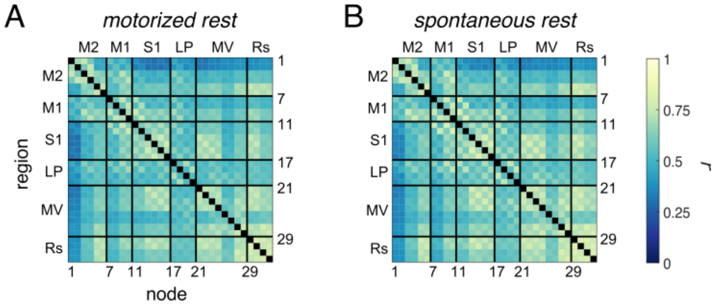


**Supplementary Figure 3. Average correlations during rest in the spontaneous locomotion condition.**

**A**. The correlation between node pairs during rest in the spontaneous locomotion condition, averaged across mice. **B**. The correlations shown in A with the Fisher transformation applied.

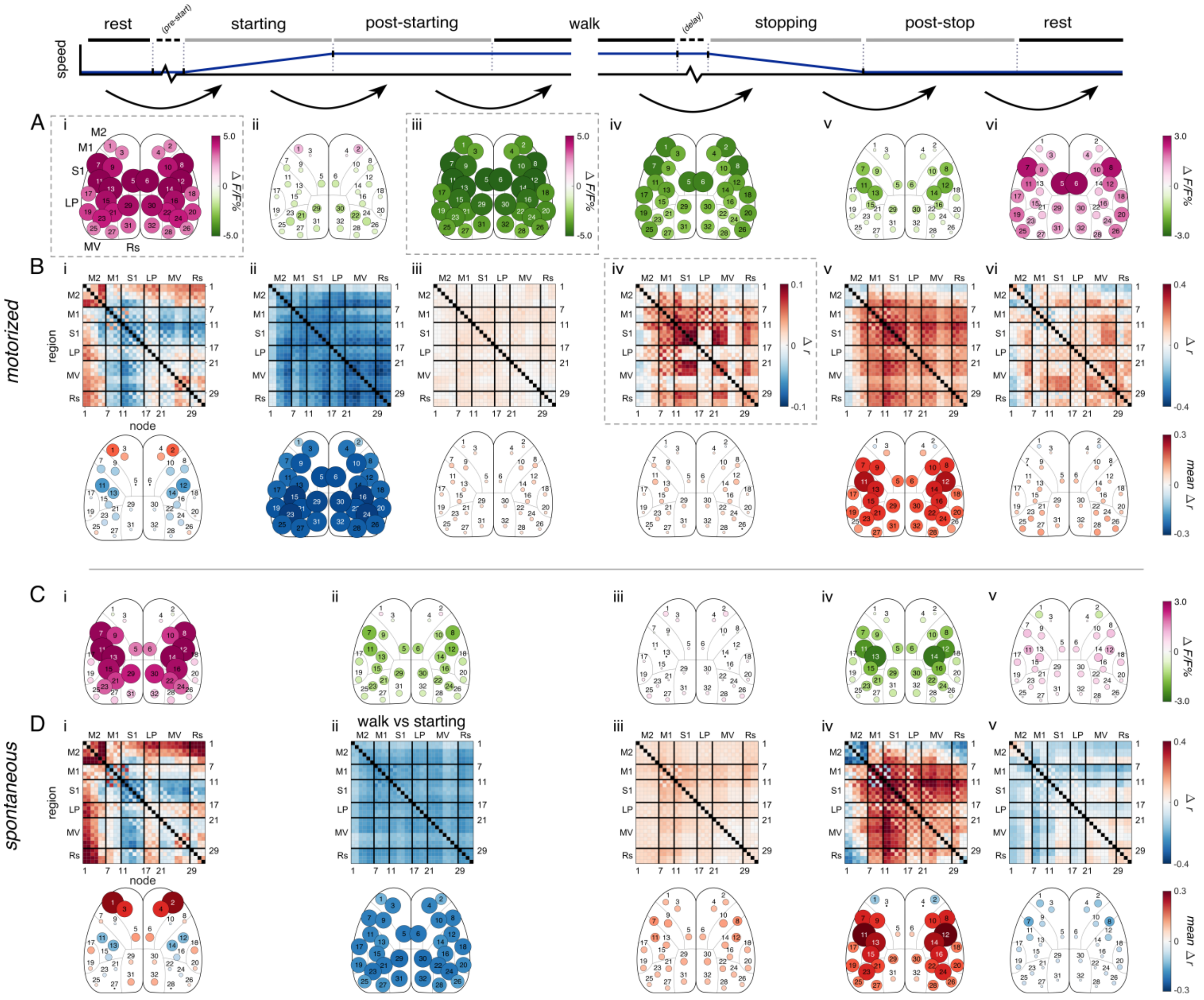


**Supplementary Figure 4. Change in correlation across sequential behavior periods.**

**A & B.** Significant changes in fluorescence (A) and correlations (B) across sequential behavior periods in the motorized treadmill condition, calculated and displayed as in Figures 2 & 3. Directly above each correlation matrix, the black arrow indicates the adjacent behavior periods being compared (i-vi). Positive values indicate increases in fluorescence or correlation across the change in behavior period. Delay periods were not included here. **C & D.** Similar to A & B, but for the spontaneous locomotion condition. Note that the spontaneous condition has no post-starting period, so the change from starting to walk (Cii & Dii) is provided instead.

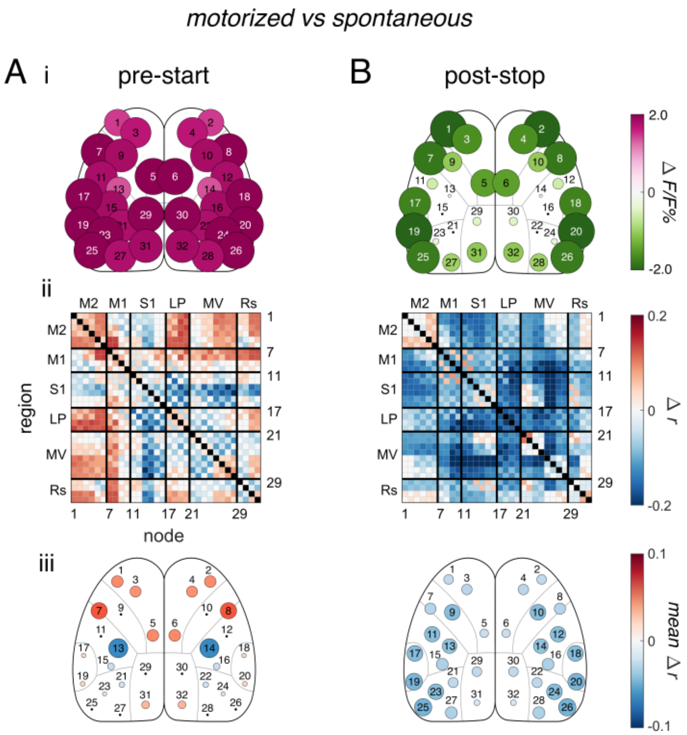


**Supplementary Figure 5. Direct comparisons of pre-start and post-stop periods across conditions.**

**A**. Significant difference in pre-start period fluorescence and FC between motorized and spontaneous locomotion conditions (motorized minus spontaneous), calculated and displayed as in Figure 3. **B**. Similar to A, but for post-stop periods.

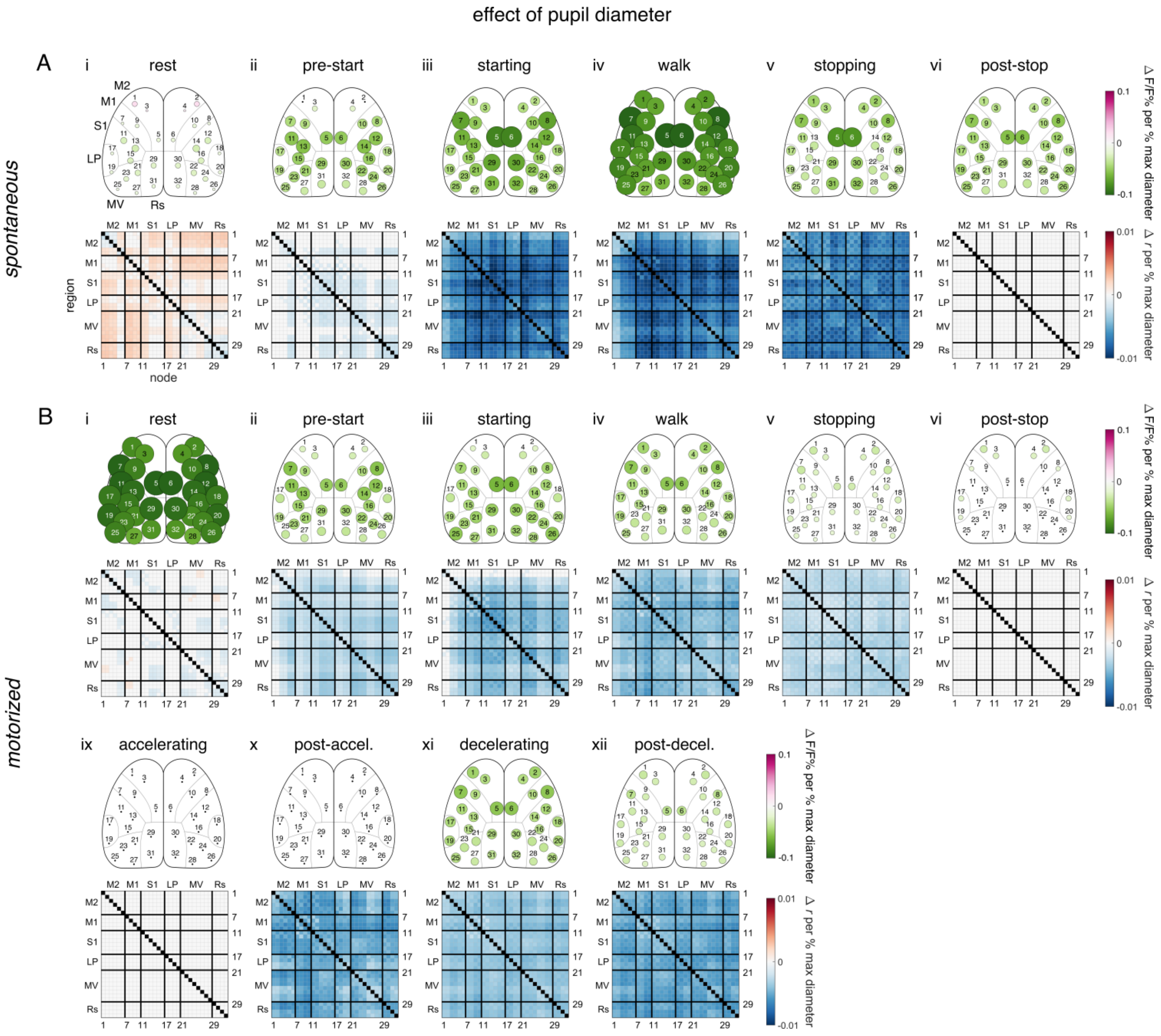


**Supplementary Figure 6. Effect of pupil diameter on behavior periods.**

**A**. Significant changes in fluorescence and correlation per % of max pupil diameter for spontaneous periods (α<0.05, permutation test with false discovery rate correction). **B**. Similar to A, but for motorized periods. Data in all panels are calculated and displayed as in Figure 2.
